# ERGA-BGE Reference Genome of *Gluvia dorsalis*: An Endemic Sun Spider from Iberian Arid Regions

**DOI:** 10.1101/2024.12.02.626363

**Authors:** Jesus Lozano-Fernandez, Marc Domènech, Attila Ibos, Thomas Marcussen, Torsten H. Struck, Rebekah Oomen, Astrid Böhne, Rita Monteiro, Laura Aguilera, Marta Gut, Francisco Câmara Ferreira, Fernando Cruz, Jèssica Gómez-Garrido, Tyler S. Alioto, Diego De Panis

**Affiliations:** Departament de Genètica, Microbiologia i Estadística, Facultat de Biologia, Universitat de Barcelona (UB), Av. Diagonal 645, 08028 Barcelona, Catalonia, Spain; Institut de Recerca de la Biodiversitat (IRBio), Universitat de Barcelona (UB), Av. Diagonal 645, 08028 Barcelona, Catalonia, Spain; La Prenyanosa, Lleida, Spain; Natural History Museum, University of Oslo, P.O. Box 1172, Blindern, 0318, Oslo, Norway; Centre for Ecological & Evolutionary Synthesis, University of Oslo, Blindernveien 31, 0371, Oslo, Norway; Department of Biological Sciences, University of New Brunswick Saint John, 100 Tucker Park Road, E2K5E2, Saint John, Canada; Tjärnö Marine Laboratory, University of Gothenburg, Hättebäcksvägen 7, Gothenburg, 45296, Sweden; Centre for Coastal Research, University of Agder, Universitetsveien 25, 4630, Kristiansand, Norway; Leibniz Institute for the Analysis of Biodiversity Change, Museum Koenig Bonn, Adenauerallee 127, 53113 Bonn, Germany; Centro Nacional de Análisis Genómico (CNAG), C/Baldiri Reixac, 4, 08028 Barcelona, Spain; Universitat de Barcelona (UB), Barcelona, Spain; Leibniz Institute for Zoo and Wildlife Research, Alfred-Kowalke-Straße 17, 10315 Berlin, Germany; Berlin Center for Genomics in Biodiversity Research (BeGenDiv), Königin-Luise-Straße 6-8, 14195 Berlin, Germany

**Keywords:** *Gluvia dorsalis*, genome assembly, European Reference Genome Atlas, Biodiversity Genomics Europe, Earth Biogenome Project, Arachnida, Solifugae, Daesiidae, *Araña camello ibérica*, *Aranya camell ibèrica*

## Abstract

The reference genome of *Gluvia dorsalis* is the first of its order Solifugae (sun spiders), offering insights into adaptations to arid environments and the evolutionary history of arachnids. The entirety of the genome sequence was assembled into 5 contiguous chromosomal pseudomolecules. This chromosome-level assembly encompasses 787 Mb, composed of 51 contigs and 10 scaffolds (including the mitogenome), with contig and scaffold N50 values of 38 Mb and 199 Mb, respectively.

## Introduction

*Gluvia dorsalis* (Latreille, 1817) is a member of the Daesiidae family within the arachnid order Solifugae. Members of this group, commonly known as sun spiders or camel spiders, inhabit arid environments, particularly warm deserts with sparse vegetation, and are rarely found in Europe. Only two species of sun spiders are known to be present in Western Europe: *Gluvia dorsalis*, endemic to the arid regions of Spain and Portugal, and *G. brunnea* Pertegal, Barranco, De Mas, and Moya-Laraño, 2024, recently described in a small region of southern Spain (Pertegal et al., 2024). *Gluvia dorsalis* is a ground-dwelling arachnid that can reach between 15 and 22 mm in length, with females being larger than males. Although not venomous, it is a fast-moving nocturnal predator that usually hides under stones during daytime. It has a yellow, orange, or reddish prosoma and legs, and a dark abdomen. The two pedipalps are highly developed, and they bear a membranous suctorial organ at the tips that allows the sun spider to capture prey and climb smooth surfaces. The diet of *G. dorsalis* includes mainly ants and spiders (Hrušková-Martišová et al., 2010), although it can potentially consume a wider range of prey. Sun spiders possess powerful pincer-like chelicerae projected forward that allow them to capture and consume large prey. *Gluvia dorsalis* can be distinguished from its relative *G. brunnea* mainly by its coloration. While *G. dorsalis* has yellow areas in the palps and legs, *G. brunnea* is dorsally completely brown. In addition, mature individuals of *G. brunnea* bear a hypertrophied seta on the basal and internal part of coxa, which is absent in *G. dorsalis*. These recent findings indicate the potential for greater genetic diversity in sun spiders than previously assumed, though further investigation is needed to confirm this interpretation.

Developing a high-quality reference genome for *G. dorsalis* is crucial for two reasons. Firstly, this information will help to improve our understanding of genomic adaptations to extreme environments, in particular to extremely hot and dry regions. Moreover, gaining a better knowledge of the genomics of this sun spider is relevant to understanding the distribution patterns of this species in particular, as well as the global distribution of sun spiders in general. Secondly, evolutionary relationships within arachnids are still among the most challenging phylogenetic relationships to resolve within animals (Lozano-Fernandez et al., 2019; Ballesteros et al., 2022). This is due to the old origin of the group, the rapid radiation of all their orders, and the multiple Whole Genome Duplication (WGD) events that some of these groups have undergone, some independent and some shared between orders (Leite et al., 2018). In the present study, we present the first genome at the chromosome level for any species from the order Solifugae, which will allow us to test whether this group of arachnids has undergone WGD. Moreover, it will greatly help to locate the position of this group, still poorly represented in genetic databases, within the arachnid tree of life.

The generation of this reference resource was coordinated by the European Reference Genome Atlas (ERGA) initiative’s Biodiversity Genomics Europe (BGE) project, supporting ERGA’s aims of promoting transnational cooperation to promote advances in the application of genomics technologies to protect and restore biodiversity (Mazzoni et al., 2023). This species falls within the regional reach of the Catalan Initiative for the Earth BioGenome Project (CBP), which is linked to ERGA (Corominas et al., 2024).

## Materials & Methods

ERGA’s sequencing strategy includes Oxford Nanopore Technology (ONT) and/or Pacific Biosciences (PacBio) for long-read sequencing, along with Hi-C sequencing for chromosomal architecture, Illumina Paired-End (PE) for polishing (i.e. recommended for ONT-only assemblies), and RNA sequencing for transcriptomic profiling, to facilitate genome assembly and annotation.

### Sample and Sampling Information

On August 1^st^, 2023, an adult individual of *G. dorsalis* (sex undetermined, however, morphological appearance suggested a female specimen) was sampled by Attila Ibos. The species was first identified morphologically by Marc Domènech and confirmed through COI barcoding by Jesus Lozano-Fernandez from Universitat de Barcelona. The specimen was caught directly with a plastic tube from the ground in Prenyanosa, Lleida (Catalonia, Spain). Sampling was conducted under the permit SF/0117/23, issued by the Catalan Government. The specimen’s tissues (e.g.: cephalothorax, abdomen, and legs) were snap-frozen immediately after harvesting and stored in liquid nitrogen until DNA extraction.

### Vouchering information

Frozen reference tissue material from the sequenced individual (Figure 1) is available at the Biobank of the Museo Nacional de Ciencias Naturales in Madrid (Spain) under the voucher ID MNCN-ADN 151.722. Physical reference material from another individual of the same population has been deposited in the same museum under the accession number MNCN20.02/22140.

**Figure 1.**
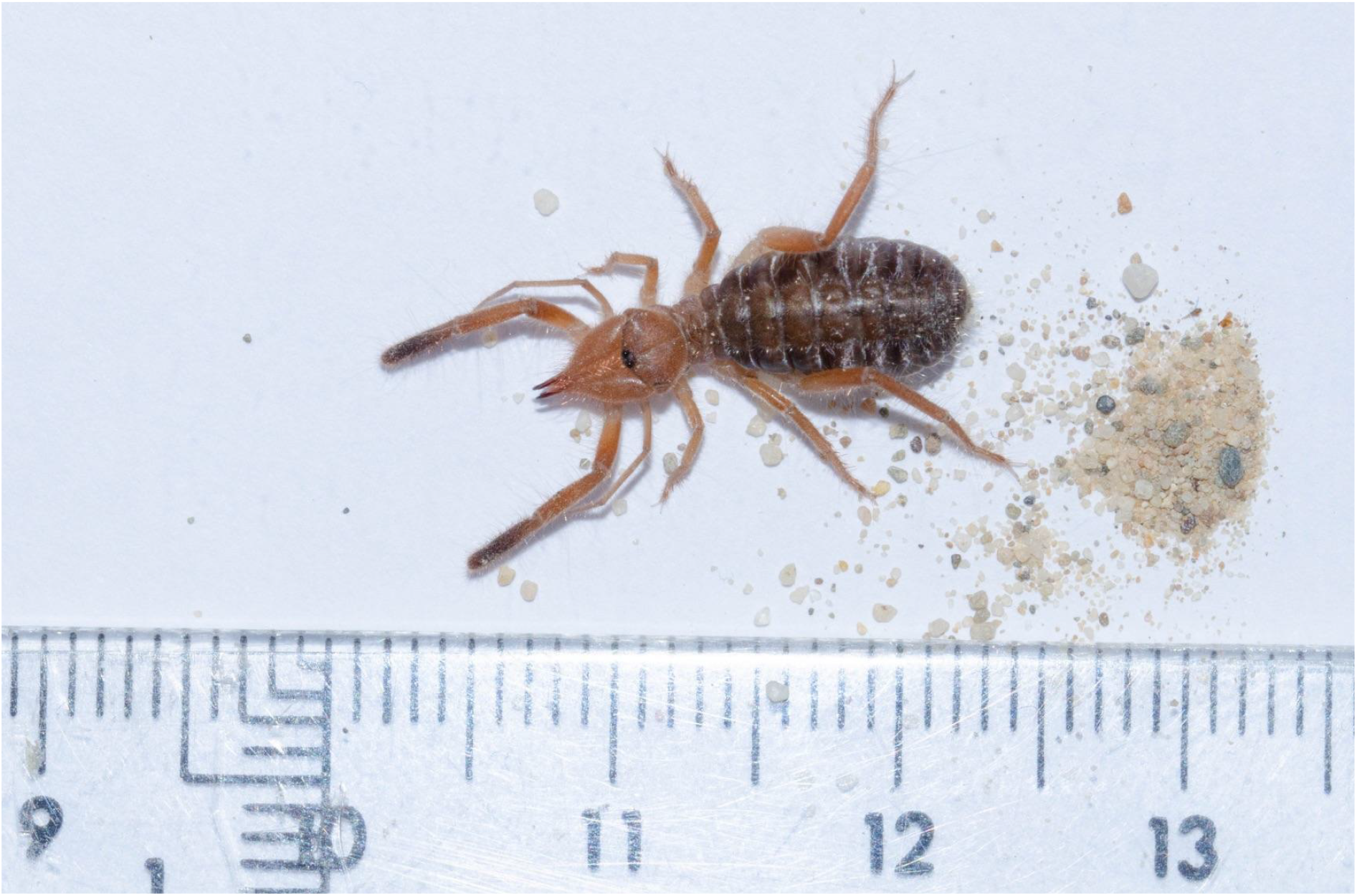
Electronic voucher image of the sequenced individual of *Gluvia dorsalis*. The image, along with two others, is available in ERGA’s EBI BioImageArchive dataset (www.ebi.ac.uk/biostudies/bioimages/studies/S-BIAD1012?query=ERGA) under accession ID SAMEA114558555.

### Data Availability

*Gluvia dorsalis* and the related genomic study were assigned to Tree of Life identifier (ToLID) ‘qqGluDors1’ and all sample, sequence, and assembly information are available under the umbrella BioProject PRJEB76507. The sample information is available at the following BioSample accessions: SAMEA115728209, SAMEA114558560, and SAMEA114558561.

The genome assembly is accessible from ENA under accession number GCA_964187665.1 and the annotated genome is available through the Ensembl Rapid Release page (projects.ensembl.org/erga-bge). Sequencing data produced as part of this project are available from ENA at the following accessions: ERX13168338, ERX12623869, ERX13168339, and ERX12623871. Documentation related to the genome assembly and curation can be found in the ERGA Assembly Report (EAR) document available at github.com/ERGA-consortium/EARs/tree/main/Assembly_Reports/Gluvia_dorsalis/qqGluDors1.

Further details and data about the project are hosted on the ERGA portal at portal.erga-biodiversity.eu/organism/SAMEA114558555.

### Genetic Information

The estimated genome size, based on ancestral taxa is 1.1 Gb, while the estimation based on reads kmer profiling is 0.79 Gb. This is a diploid genome with a haploid number of 5 chromosomes (2n=10). Information for this species was retrieved from Genomes on a Tree (Challis et al., 2023).

### DNA/RNA processing

DNA was extracted from the cephalothorax and abdomen using the Blood & Cell Culture DNA Mini Kit (Qiagen) following the manufacturer’s instructions. DNA quantification was performed using a Qubit dsDNA BR Assay Kit (Thermo Fisher Scientific), and DNA integrity was assessed using a Femtopulse system (Genomic DNA 165 Kb Kit, Agilent). DNA was stored at 4ºC until use.

RNA was extracted using an RNeasy Mini Kit (Qiagen) according to the manufacturer’s instructions. RNA was extracted from two different specimen body parts: leg and cephalothorax. RNA quantification was performed using the Qubit RNA BR Kit and RNA integrity was assessed using a Bioanalyzer 2100 system (Eukaryote Total RNA Pico Kit, Agilent). RNA was pooled in a 1:5 (leg:cephalothorax) ratio before library preparation and stored at -80ºC until use.

### Library Preparation and Sequencing

A long-read whole genome library was prepared using the SQK-LSK114 kit and sequenced on a PromethION P24 A series instrument (Oxford Nanopore Technologies). For short-read whole genome sequencing (WGS), a library was constructed with the KAPA Hyper Prep Kit (Roche) for subsequent sequencing on an Illumina platform. Hi-C library preparation, using cephalothorax and leg tissue, was conducted with the ARIMA High Coverage Hi-C Kit (Arima) and further processed with the KAPA Hyper Prep Kit for Illumina sequencing (Roche). The RNA library, generated from the pooled sample, was prepared with the KAPA mRNA Hyper Prep Kit (Roche). All short-read libraries were sequenced on the Illumina NovaSeq 6000 instrument. In total, 116x Oxford Nanopore, 102x Illumina WGS shotgun, and 97x HiC data were sequenced to generate the assembly.

### Genome Assembly Methods

The genome was assembled using the CNAG CLAWS pipeline (Gomez-Garrido, 2024). Briefly, reads were preprocessed for quality and length using Trim Galore v0.6.7 and Filtlong v0.2.1, and initial contigs were assembled using NextDenovo v2.5.0, followed by polishing of the assembled contigs using HyPo v1.0.3, removal of retained haplotigs using purge-dups v1.2.6 and scaffolding with YaHS v1.2a.

Finally, assembled scaffolds were curated via manual inspection using Pretext v0.2.5 with the Rapid Curation Toolkit (gitlab.com/wtsi-grit/rapid-curation) to remove any false joins and incorporate any sequences not automatically scaffolded into their respective locations in the chromosomal pseudomolecules (or super-scaffolds). The mitochondrial genome was assembled as a single circular contig of 14,734 bp using the FOAM pipeline v0.5 (github.com/cnag-aat/FOAM) and included in the released assembly (GCA_964187665.1). Summary analysis of the released assembly was performed using the ERGA-BGE Genome Report ASM Galaxy workflow (De Panis, 2024), incorporating tools such as BUSCO v5.5, Merqury v1.3, and others (see reference for the full list of tools).

## Results

### Genome Assembly

The genome assembly had a total length of 787,034,199 bp in 10 scaffolds including the mitogenome (Figures 2 and 3), with a GC content of 39.73%. It featured a contig N50 of 37,604,012 bp (L50=8) and a scaffold N50 of 198,509,873 bp (L50=2). There were 41 gaps, totaling 8,200 kb in cumulative size. The single-copy gene content analysis using the Arachnida database with BUSCO resulted in 94.7% completeness (93.3% single and 1.4% duplicated). Additionally, 95.6% of reads k-mers were present in the assembly and the assembly has a base accuracy Quality Value (QV) of 48.05 as calculated by Merqury.

**Figure 2.**
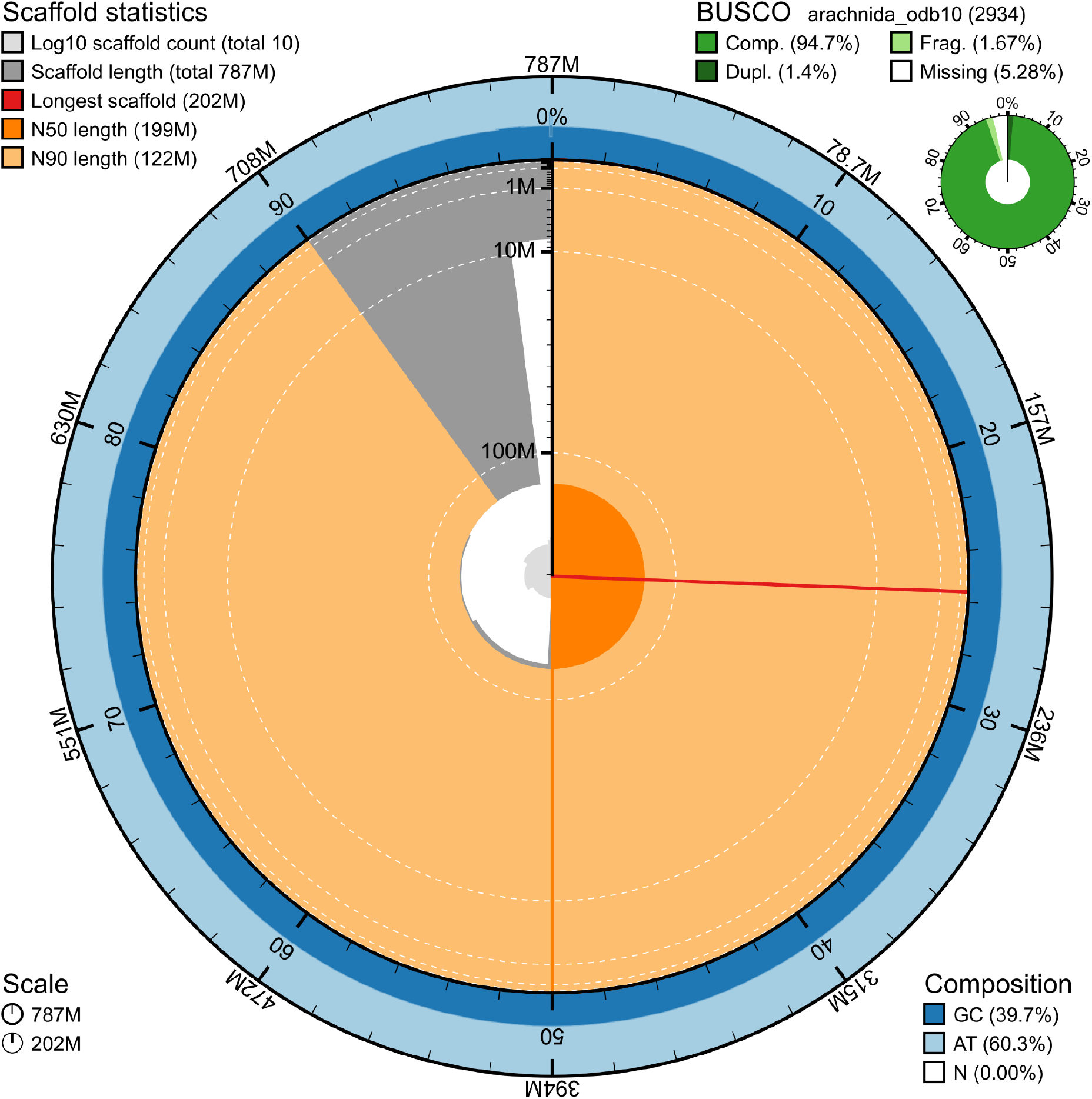
Snail plot summary of assembly statistics. The main plot is divided into 1,000 size-ordered bins around the circumference, with each bin representing 0.1% of the 787,034,199 bp assembly including the mitochondrial genome. The distribution of sequence lengths is shown in dark grey, with the plot radius scaled to the longest sequence present in the assembly (201,641,468 bp, shown in red). Orange and pale-orange arcs show the scaffold N50 and N90 sequence lengths (198,509,873 and 122,092,752 bp), respectively. The pale grey spiral shows the cumulative sequence count on a log-scale, with white scale lines showing successive orders of magnitude. The blue and pale-blue area around the outside of the plot shows the distribution of GC, AT, and N percentages in the same bins as the inner plot. A summary of complete, fragmented, duplicated, and missing BUSCO genes found in the assembled genome from the Arachnida database (odb10) is shown on the top right.

**Figure 3.**
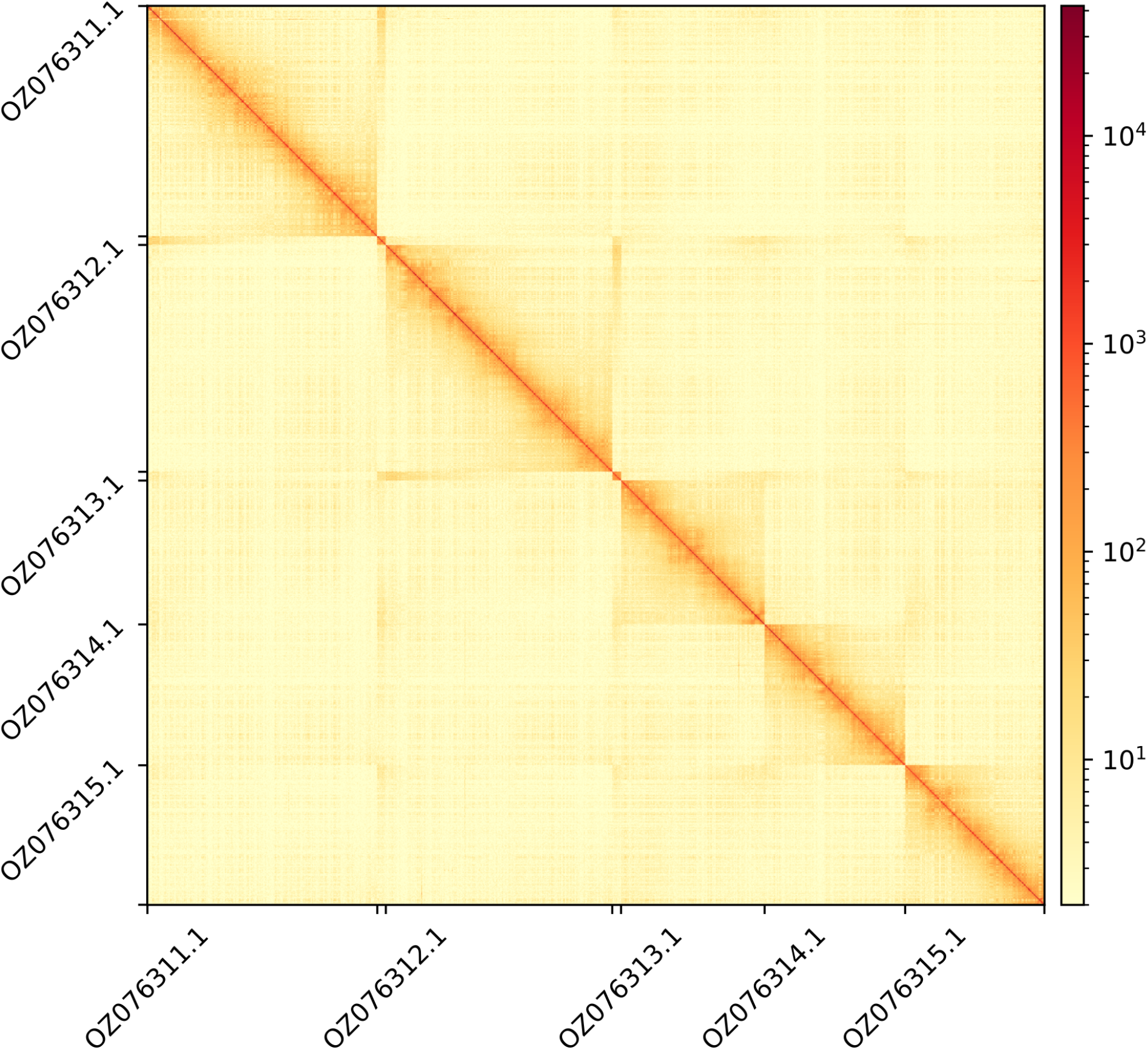
Hi-C contact map showing spatial interactions between regions of the genome. The diagonal corresponds to intra-chromosomal contacts, depicting chromosome boundaries. The frequency of contacts is shown on a logarithmic heatmap scale. Hi-C matrix bins were merged into a 200 kb bin size for plotting. From the 10 Scaffolds including the mitogenome, only the GenBank names of the five chromosomes are shown.

## Acknowledgements

We thank Alberto Narro for his willingness to help by providing samples from other populations. We acknowledge the support of the Freiburg Galaxy Team: Saim Momin and Björn Grüning, Bioinformatics, University of Freiburg (Germany), funded by the German Federal Ministry of Education and Research BMBF grant 031 A538A de.NBI-RBC and the Ministry of Science, Research and the Arts Baden-Württemberg (MWK) within the framework of LIBIS/de.NBI Freiburg. We would like to acknowledge the assembly reviewer, Tom Mathers, from the Wellcome Sanger Institute.

## Author Contributions

JLF coordinated the project, AI and MD collected the species, MD and JLF identified the species, JLF and MD sampled and preserved biological material and provided metadata, RM, TM, RO, THS, and AsB provided sampling and metadata support and management, LA and MG extracted DNA, prepared libraries, and performed sequencing, FCF, JGG and FC performed genome assembly and curation under the supervision of TSA, DDP generated the analysis and report. All authors contributed to the writing, review, and editing of this genome note and read and approved the final version.

## Literature Cited

Ballesteros, J. A., Santibáñez-López, C. E., Baker, C. M., Benavides, L. R., Cunha, T. J., Gainett, G., Ontano, A. Z., Setton, E. V. W., Arango, C. P., Gavish-Regev, E., Harvey, M. S., Wheeler, W. C., Hormiga, G., Giribet, G., & Sharma, P. P. (2022). Comprehensive Species Sampling and Sophisticated Algorithmic Approaches Refute the Monophyly of Arachnida. Molecular Biology and Evolution, 39(2). 10.1093/molbev/msac021

Challis, R., Kumar, S., Sotero-Caio, C., Brown, M., & Blaxter, M. (2023). Genomes on a Tree (GoaT): A versatile, scalable search engine for genomic and sequencing project metadata across the eukaryotic tree of life. Wellcome Open Research, 8, 24. 10.12688/wellcomeopenres.18658.1

Corominas, M., Marquès-Bonet, T., Arnedo, M. A., Bayés, M., Belmonte, J., Escrivà, H., Fernández, R., Gabaldón, T., Garnatje, T., Germain, J., Niell, M., Palero, F., Pons, J., Puigdomènech, P., Initiative For The Earth BioGenome Project, T. C., Catalan initiative for the Earth BioGenome Project, Arroyo, V., Cuevas-Caballé, C., Obiol, J. F., … Guigó, R. (2024). The Catalan initiative for the Earth BioGenome Project: Contributing local data to global biodiversity genomics. NAR Genomics and Bioinformatics, 6(3), qae075. 10.1093/nargab/lqae075

De Panis, D. (2024). ERGA-BGE Genome Report ASM analyses (one-asm WGS Illumina PE + HiC). WorkflowHub. 10.48546/WORKFLOWHUB.WORKFLOW.1103.2

Gomez-Garrido, J. (2024). CLAWS (CNAG’s long-read assembly workflow in Snakemake). WorkflowHub. 10.48546/WORKFLOWHUB.WORKFLOW.567.2

Hrušková-Martišová, M., Pekár, S., & Cardoso, P. (2010). Natural history of the Iberian solifuge Gluvia dorsalis (Solifuges: Daesiidae). The Journal of Arachnology, 38(3), 466–474. 10.1636/Hi09-104.1

Leite, D. J., Baudouin-Gonzalez, L., Iwasaki-Yokozawa, S., Lozano-Fernandez, J., Turetzek, N., Akiyama-Oda, Y., Prpic, N.-M., Pisani, D., Oda, H., Sharma, P. P., & McGregor, A. P. (2018). Homeobox Gene Duplication and Divergence in Arachnids. Molecular Biology and Evolution, 35(9), 2240–2253. 10.1093/molbev/msy125

Lozano-Fernandez, J., Tanner, A. R., Giacomelli, M., Carton, R., Vinther, J., Edgecombe, G. D., & Pisani, D. (2019). Author Correction: Increasing species sampling in chelicerate genomic-scale datasets provides support for monophyly of Acari and Arachnida. Nature Communications, 10, 4534. 10.1038/s41467-019-12259-6

Mazzoni, C., Ciofi, C., & Waterhouse, R. (2023). Biodiversity: An atlas of European reference genomes. Nature, 619, 252–252. 10.1038/d41586-023-02229-w

Pertegal, C., Barranco, P., De Mas, E., & Moya-Laraño, J. (2024). More Than 200 Years Later: Gluvia brunnea sp. nov. (Solifugae, Daesiidae), a Second Species of Camel Spider from the Iberian Peninsula. Insects, 15(4), 284. 10.3390/insects15040284

